# A causative SNP in the promoter of myogenin is essential for myogenic differentiation

**DOI:** 10.1101/2024.08.01.606143

**Authors:** Zhuhu Lin, Xiaoyu Wang, Ziyun Liang, Rong Xu, Meilin Chen, Xian Tong, Chenggan Li, Yanyun Xiong, Renqiang Yuan, Yaosheng Chen, Yunxiang Zhao, Xiaohong Liu, Delin Mo

**Author notes:** **Corresponding author:** Delin Mo,; Xiaohong Liu,; Yunxiang Zhao,. **Author information:** Zhuhu Lin,; Xiaoyu Wang,; Ziyun Liang,; Rong Xu,; Meilin Chen,; Xian Tong,; Chenggan Li,; Yanyun Xiong,; Renqiang Yuan, Yaosheng Chen,.

## Abstract

Single nucleotide polymorphisms (SNPs) widely existing in different breeds genome represent population-specific. Under the influence of long-term evolution and artificial selection, there are a large number of SNPs between western lean-type pig breeds and Chinese indigenous pig breeds, but until now, little is known about their roles in inter-breed differences. Our study revealed SNP rs3471653254 C>T generated from the two types of pigs mentioned above, located in the promoter shared by MyoG and Myoparr, played an important role in the differentiation of myoblast by influencing the enrichment of HOXA5 to regulate the transcription of MyoG and Myoparr. Meanwhile, Myoparr could be used as the sponge of mir-30b-3p which repressed myogenic differentiation and muscle regeneration through targeting MyoD. Our results indicated that SNP rs3471653254 C>T is essential for myogenic differentiation and regeneration and could be used as an ideal site for increasing lean meat production in pigs.

## Introduction

The skeletal muscle, representing nearly 40% of adult tissue in mammal, is the most abundant and highly complex tissue(Bentzinger et al. 2012; Frontera and Ochala 2015). The development of vertebrate skeletal muscle includes the specification of mesodermal precursor cells into myoblasts, followed by subsequent proliferation, differentiation and fusion of these cells into multinucleated myotubes during embryonic development, continually regenerating and self-renewal throughout life(Molkentin and Olson 1996). Myogenesis is a highly ordered process regulated spatiotemporally by a lot of regulatory networks and signal pathways(Braun and Gautel 2011). Myogenic regulatory factors (MRFs), including Myf5, MyoD, Myogenin (MyoG) and MRF4, are the core components regulating myogenesis. In early myogenesis, Myf5, MyoD are in charge of mediating the initial specification of skeletal myoblasts. MyoG, MRF4, and myocyte-specific enhancer factors (MEF2A and MEF2C) induce differentiation of these specified cells(Nie et al. 2020; Zhu et al. 2021). Myosin heavy chain (MyHC) protein, also greatly functions in muscle contraction(Forcales et al. 2012; Blum and Dynlacht 2013; Buckingham and Rigby 2014).

In addition to the myogenic transcription factors, non-coding RNAs such as microRNAs (miRNAs), long non-coding RNAs (lncRNAs) and circular RNAs (circRNAs) have been reported to be involved in the development of skeletal muscle and play critical roles in myogenesis(Zhang et al. 2018; Gorospe et al. 2020; Liu et al. 2020). LncRNAs are pervasively transcribed polyadenylated RNA molecules that are >200 bp in length(Michalik et al. 2014; Lee et al. 2016). While a striking 75% of human genome was transcribed, only 2% of these genes encode proteins, and the vast majority of transcription produces lncRNAs, as reported by the ENCODE Project Consortium(Consortium 2012). Increasing evidence indicates that promoter-associated lncRNAs act in cis to regulate the transcription of protein-coding genes, particularly as developmental regulators(Hamazaki et al. 2015). To give a prominent example, one of its functional mechanisms is provided by competing endogenous RNA (ceRNA), which bears miRNA binding sites. Then miRNAs would induce the decay of mRNA and translational suppression through the interaction with the complementary sequences in the 3’UTR of target gene(Bartel 2009; Akgul and Erdogan 2018), then enabling them to have important regulatory roles in diverse physiological and developmental processes(Friedman et al. 2009).

Single nucleotide polymorphism (SNP), is a kind of DNA sequence polymorphism caused by a single nucleotide variation at the genomic level(Brookes 1999). On the other hand, nucleotide mutation is an irreversible sequence variation in DNA, which in essence includes all variations that occur in the species genome either spontaneously or non-spontaneously(Brown 2002). The distribution of SNPs is population-specific, which may be one of the reasons for phenotypic differences among populations. Under the influence of long-term evolution and artificial selection, pig breeds from all over the world show differences in meat production, growth rate, feed conversion ratio etc(Wang et al. 2022). Western lean-type breeds, such as Duroc pigs (DR), have been intensively selected over the past several decades for high muscle mass, while Chinese indigenous breeds, such as Guangdong small-ear spotted pig (GS), have lower growth rates and lean meat percentage without sustained artificially selection(Suzuki et al. 1991; Tang et al. 2007). Such a significant difference in meat production traits must be caused by genetic variation. Moreover, SNPs are widely distributed in the swine genome, with an average distribution interval of 300 to 400 bp(Jungerius et al. 2005). By comparing the genomes, it was found that there are a great quantity of SNPs between western lean-type pig breeds and Chinese indigenous pig breeds(Amaral et al. 2009; Kerstens et al. 2009; Ramos et al. 2009). Until now, several studies have indicated that SNPs are implicated in skeletal muscle formation, but little has been known about its roles and molecular mechanism on inter-breed differences(Tong et al. 2015; Khanal et al. 2021; Reguero et al. 2021; Ma et al. 2022). What’s more, significant phenotypic variations occurred in Yuedong Black pigs (YDB) from 1975 to 2006. At present, YDB assume a new body-weight and body-shape, unexpectedly close to western lean-type pigs. By whole genome re-sequencing of Large White, Landrace, Duroc and 16 kinds of Chinese indigenous pig breeds (including YDB), we found that the MyoG gene sequence of YDB is much closer to western lean-type pigs. Further, gene introgression analysis revealed that MyoG gene of YDB originated from Duroc(Wang 2021). Based on the previous research, we hypothesize that the population-specific SNPs of myogenin sequence lead to the body change of YDB, but also serve as one of the main reasons for the difference in meat production between the two kinds of pig breeds. Therefore, we conduct research on SNPs among lean-type pig breeds and Chinese indigenous, providing theoretical knowledge support for explaining the differences of two kinds of pig breeds. Proceed to the next step, these SNPs could be expanded on techniques such as marker-assisted selection or gene editing for increasing lean meat production. In this research, SNP rs3471653254 C>T, located in the promoter at 299 base pairs before translation start site (TSS) of MyoG, played an important role in myoblast differentiation. It regulates the transcription of MyoG and Myoparr by influencing the enrichment of transcription factor homeobox A5 (HOXA5) on the shared promoter. Meanwhile, Myoparr can be used as the sponge of mir-30b-3p, which constitutes a ceRNA. We further found mir-30b-3p overexpression represses myogenesis in C2C12 cells and muscle regeneration along with attenuating myogenic genes expression, whereas mir-30b-3p depletion has the opposite effects. Furthermore, we demonstrate mir-30b-3p regulates myoblast differentiation by targeting MyoD. In a word, SNP rs3471653254 C>T was found to be a vital and multifunctional gene site regulating myogenesis. Moreover, our results elucidate a novel myogenic inhibitor mir-30b-3p and its regulatory network in myogenesis.

## Result

### SNPs identified among western lean-type pig breeds and Chinese indigenous pig regulate myoblast differentiation

Yuedong black pig, one of the indigenous pig breeds in the south of China, has undergone significant changes in weight and body shape compared to 30 years ago. Gene introgression analysis revealed that a long DNA fragment of approximately 950kb including myogenin gene originated from Duroc, which is line with its history of introducing hybridization. By comparing the genome sequences of myogenin between 4 western lean-type pig breeds and 16 Chinese indigenous pig breeds, a total of 11 SNPs that maintaining stable difference between the two kinds of pig breeds were selected. Then two kinds of MyoG expression plasmid were constructed according to SNPs based on western lean-type pig breed (DR) and Chinese indigenous pig breed (GS) (Figure 1A). The whole MyoG was overexpressed in porcine primary myoblasts and 293T cells using plasmid transfection (Figure 1B-1D and Figure 1L-1M). As a result, the cells transfected with MyoG plasmid of DR show remarkably higher level of myogenic differentiation as determined by immunofluorescence staining of MyHC (Figure 1E). Compared with the GS plasmid, the fusion index and differentiation index of porcine primary myoblasts transfected with DR plasmid increased by nearly 13% (Figure 1E). In addition, the expression of genes related to myogenesis was monitored by qPCR and Western blot. MYHC and MyoD showed higher expression in the cells transfected with MyoG plasmid of DR. (Figure 1F-1K). In the same way for mouse specie, when the MyoG plasmid of DR overexpressed in C2C12 cells, significant promotion was observed in both myoblast differentiation and myogenic marker genes expression (Figure S1A-S1J). To explore which SNP plays the key role, the MyoG plasmids were respectively designed to mutate on the upstream or downstream (Figure S1K). The mRNA and protein expression level of MyoG shows almost no difference between downstream mutation and DR’s MyoG plasmid (Figure), but decrease significantly when mutations occurred in upstream (Figure 1N-1P). These results prove that SNPs in the upstream region of MyoG regulate the expression of myogenic genes and myoblast differentiation.

**Figure 1.**
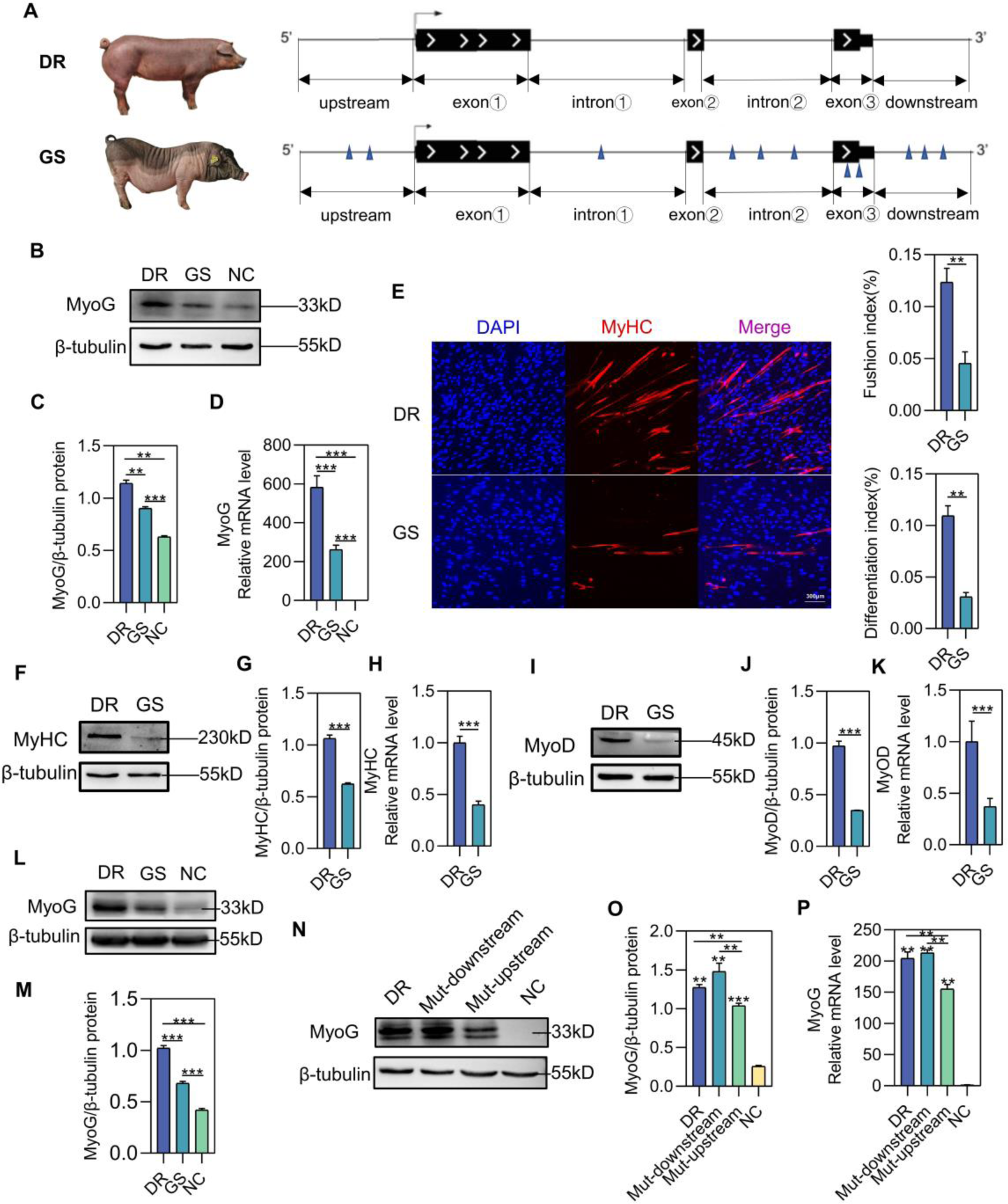
SNP regulates myoblast differentiation and the expression of myogenic genes. (A) Construction of two kinds of MyoG vectors including selected 11 SNPs that maintaining stable difference between the two pig breeds. (B-C) Western blot analysis of MyoG overexpression in porcine primary myoblasts transfected with control pcDNA3.1 or two kinds of MyoG vectors from Figure 1A after 48 h (n=3). (D) QPCR analysis of MyoG overexpression in porcine primary myoblasts transfected with control pcDNA3.1 or two kinds of MyoG vectors after 48 h (n=3). (E) Representative photographs of MyHC immunofluorescence staining in porcine primary myoblasts differentiated for 6 days after overexpression of two kinds of MyoG vectors. Statistical results of myotube fusion index (n=3). Myotube fusion index was determined as the distribution of nucleus number in total myotubes. Statistical results of differentiation index. The differentiation index was calculated as the percentage of MyHC-positive nuclei among total nuclei(n=3). (F-K) Western blot and qPCR analysis for myogenic marker genes expression from differentiated porcine primary myoblasts for 6 days after overexpression of two kinds of MyoG vectors. (L-M) Western blot analysis of MyoG overexpression in 293T cells transfected with control pcDNA3.1 or two kinds of MyoG vectors from Figure 1A after 48 h (n=3). (N-P) Western blot and qPCR analysis for MyoG expression from 293T 2 cells transfected with control pcDNA3.1 or 3 kinds of MyoG vectors from Figure S1K after 48 h (n=3). Data are represented as mean ±SD. \**P* < 0.05; \*\**P* < 0.01; \*\*\**P* < 0.001 (Student’s *t* test).

### Identification of lncRNA Myoparr and acts as a ceRNA sponging miR-30b-3p

In order to figure out why SNPs of upstream promoter can affect the expression of myogenin and myoblast differentiation, we analyzed the sequences of myogenin upstream and identified a promoter-associated lncRNA Myoparr (Figure 2A). More importantly, Myoparr shares the same promoter together with MyoG and presented significantly higher expression after transfection with plasmid of DR over plasmid of GS (Figure 2B, Figure S2A). By using an online coding potential calculator (CPC), it indicates that lncRNA Myoparr has no coding potential, with the positive control for MyoG and the noncoding control HOTAIR, a well-documented lncRNA(Figure 2C) (Hajjari and Salavaty 2015). QPCR analyses have shown that Myoparr expression was predominantly enriched in skeletal muscle tissue among the organs of 180-day-old pigs as same as MyoG and MyoD expression (Figure 2D-2F). In addition, during skeletal muscle development, the expression patterns of Myoparr were similar to the key myogenic genes, indicating its involvement in embryonic myogenesis (Figure 2K). During C2C12 differentiation, the expression level of Myoparr was peaked on DM 2d and subsequently decreased from DM 2d to DM 8d, as the same expression trend as MyoG and MyoD (Figure 2L). Considering that lncRNAs usually sponge miRNAs to modulate gene expression, an online prediction website miRBase was employed, and miR-30b-3p was screened as a candidate target (Figure 2G-2H). To verify whether miR-30b-3p can directly target Myoparr, we cloned the Myoparr DNA segment containing either the wild-type (WT) binding sites or mutant (Mut) binding sites of miR-30b-3p into luciferase reporter plasmids (Figure 2I). Dual-luciferase reporter assays showed that overexpressing miR-30b-3p could reduce the luciferase activity of Myoparr-WT, but not affect the luciferase activity of the mutant (Figure 2J), which confirmed that miR-30b-3p was sponged with the predicted target site by Myoparr. As expected, the expression trend of miR-30b-3p was contrary to Myoparr (Figure 2K-2L), indicating their strong negative correlation.

**Figure 2.**
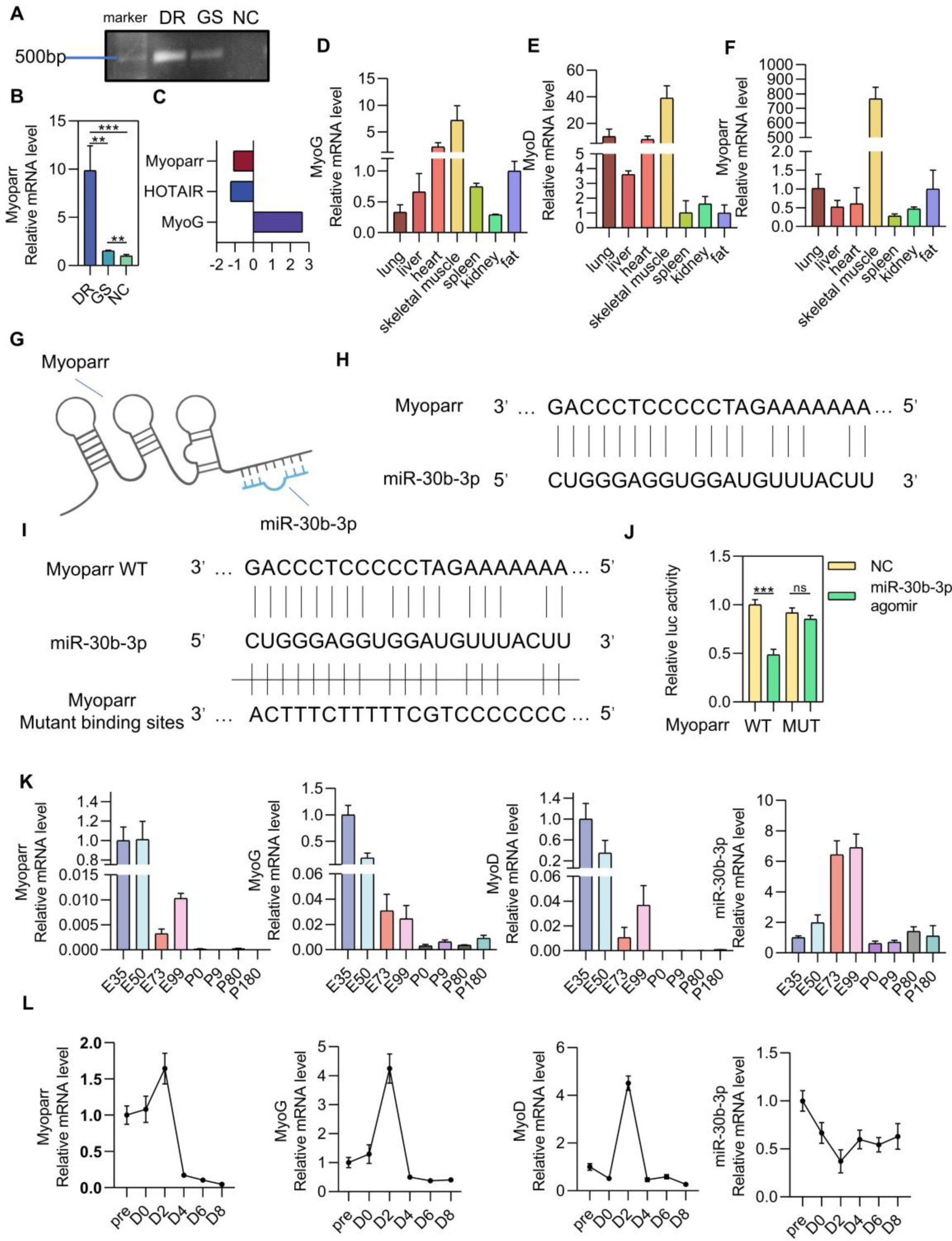
Identification of lncRNA Myoparr and validation of lncRNA Myoparr as a ceRNA sponging miR-30b-3p. (A) C2C12 cells were transfected with two kinds of MyoG vectors, pcDNA3.1 vector acted as a control. Then after 48 h PCR validated the 500bp sequence expressed by Myoparr from the extracted RNA. (B) QPCR analysis of Myoparr expression in porcine primary myoblasts transfected with control pcDNA3.1 or two kinds of MyoG vectors from Figure 1A after 48 h (n=3). (C) In silico analysis of the coding potential of Myoparr comparing with coding genes MyoG and the noncoding control HOTAIR by using CPC. (D-F) Expression pattern of MyoG, MyoD and Myoparr in various tissues in 180-day-old pigs was detected through qPCR (n=3). (G) Sketch on potential miR-30b-3p binding site in Myoparr. (H) Details of the alignment of miR-30b-3p to the binding site of Myoparr. (I) Schematic of the double luciferase assay vector with upper wildtype (WT), and lower mutant (Mut) Myoparr binding sites. (J) AgomiR-30b-3p or NC were co-transfected with psiCHECK2-Myoparr WT or psiCHECK2-Myoparr Mut vectors into 293T cells. After 48 h, Dual-Luciferase reporter assay was quantified and normalized (n=4). (K) QPCR detection of MyoD, MyoG, Myoparr and miR-30b-3p expression during pigs’ skeletal muscle development. (L) QPCR detection of MyoD, MyoG, Myoparr and miR-30b-3p expression during C2C12 cell differentiation. Data are represented as mean ± SD. \**P* < 0.05; \*\**P* < 0.01; \*\*\**P* < 0.001 (Student’s *t* test).

### MiR-30b-3p impairs myogenic differentiation

In order to investigate the function of miR-30b-3p during myogenic differentiation, porcine primary myoblasts were transfected with agomiR-30b-3p or negative control (NC) at 80% confluence and then induced to differentiation. As shown in Figure 3A, the expression of miR-30b-3p was highly up-regulated. MyHC immunofluorescence staining assay (Figure 3B) demonstrated that overexpressing miR-30b-3p prevented myognic differentiation, along with fewer myotubes and almost no thick muscle fibers (Figure 3B). Consistent with this result, the protein and mRNA expression levels of myogenic factors, including MyoD, MyoG, MyHC and Myoparr, were all decreased (Figure 3C-3L). Furthermore, the same phenomenon also appeared in C2C12 cells (Figure S3). The above data indicated that overexpression of miR-30b-3p impairs the differentiation of myoblasts. AntagomiR-30b-3p and NC were respectively transfected into porcine primary myoblasts, and the expression of miR-30b-3p was decreased nearly 70% (Figure 4A). MiR-30b-3p knockdown in primary myoblasts promoted myoblast differentiation into myotubes, evidenced by an increased number of MYHC+ cells and fusion rate with a lot of extraordinary thick myofibers in the visual field (Figure 4B). Correspondingly the expression levels of myogenic factors, such as MyoD, MyoG, MyHC and Myoparr, were markedly improved (Figure 4C-4L), which was also observed in C2C12 cells (Figure S4). The above results indicated that depletion of miR-30b-3p promotes myogenic differentiation.

**Figure 3.**
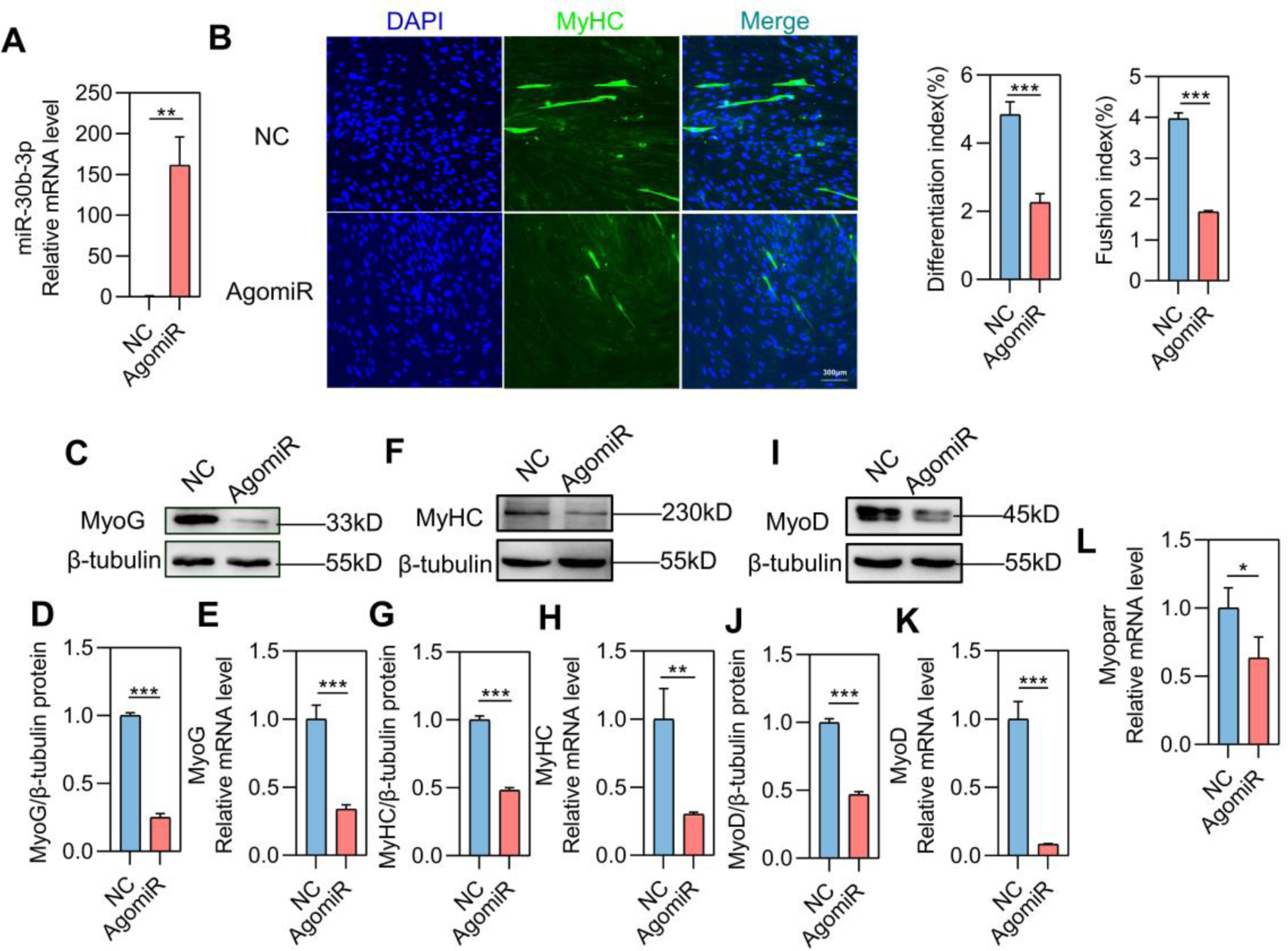
Overexpression of miR-30b-3p impairs myogenic differentiation. (A) Overexpression efficiency of agomiR-30b-3p targeting miR-30b-3p was detected in porcine primary myoblasts through qPCR (n=3). (B) Immunofluorescent stainin for MyHC in NC or agomiR-30b-3p porcine primary myoblasts was performed to detect myotube formation. Statistical results of differentiation index. The differentiation index was calculated as the percentage of MyHC-positive nuclei among total nuclei(n=3). Statistical results of myotube fusion index (n=3). Myotube fusion index was determined as the distribution of nucleus number in total myotubes. (C-L) Western blot and qPCR analysis for myogenic marker genes expression from differentiated porcine primary myoblasts for 6 days after transfected with NC or agomiR-30b-3p. Data are represented as mean ±SD. \**P* < 0.05; \*\**P* < 0.01; \*\*\**P* < 0.001 (Student’s *t* test).

**Figure 4.**
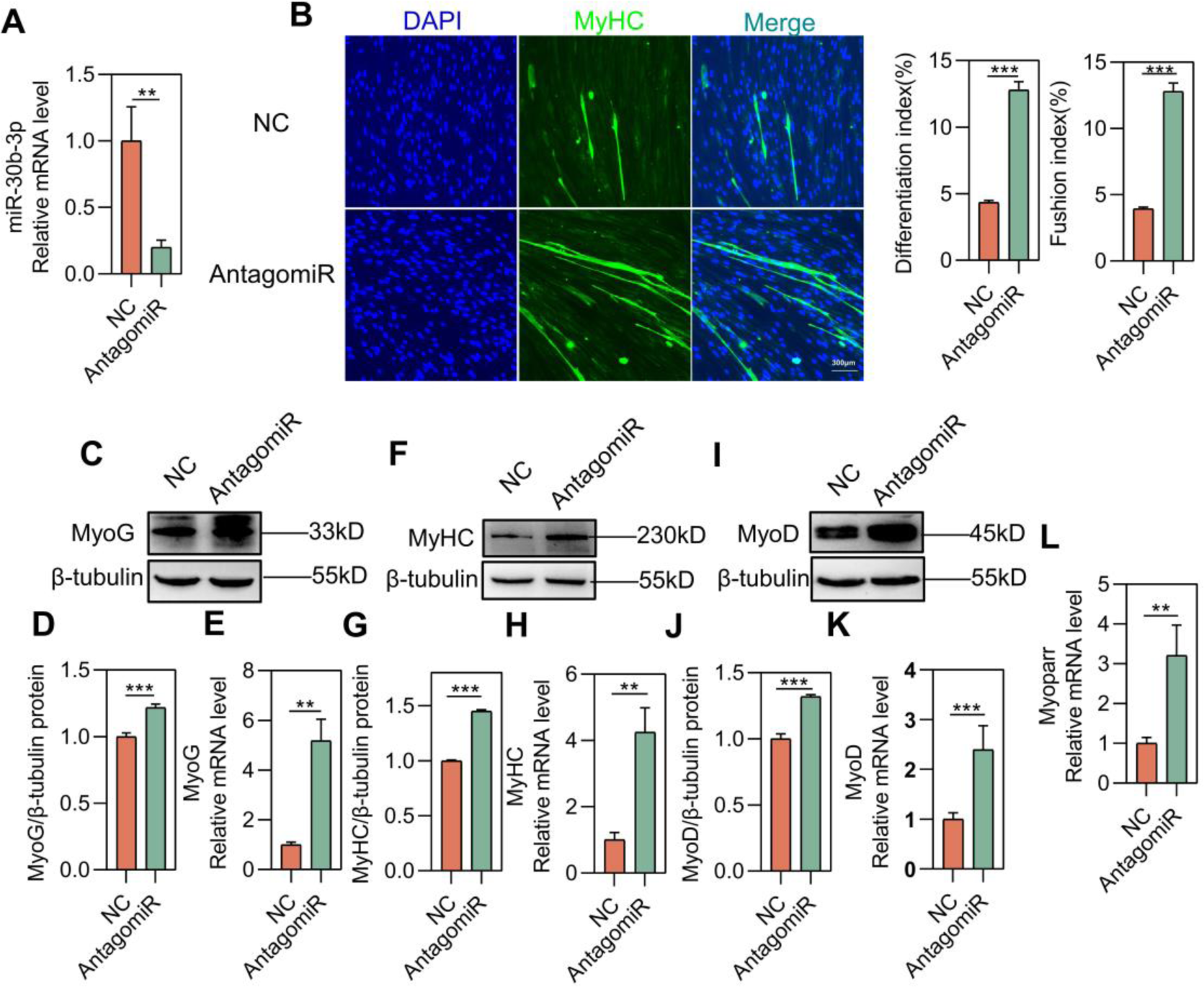
miR-30b-3p deficiency facilitates myogenic differentiation. (A) Knockdown efficiency of antagomiR-30b-3p targeting miR-30b-3p was detected in porcine primary myoblasts through qPCR (n=3). (B) Immunofluorescent stainin for MyHC in NC or antagomiR-30b-3p porcine primary myoblasts was performed to detect myotube formation. Statistical results of differentiation index. The differentiation index was calculated as the percentage of MyHC-positive nuclei among total nuclei(n=3). Statistical results of myotube fusion index (n=3). Myotube fusion index was determined as the distribution of nucleus number in total myotubes. (C-L) Western blot and qPCR analysis for myogenic marker genes expression from differentiated porcine primary myoblasts for 6 days after transfected with NC or antagomiR-30b-3p. Data are represented as mean ±SD. \**P* < 0.05; \*\**P* < 0.01; \*\*\**P* < 0.001 (Student’s *t* test).

### Elevated miR-30b-3p delays skeletal muscle regeneration

To further confirm the function of miR-30b-3p, muscle regeneration model was established, and the treatment was illustrated in Figure 5A. AgomiR-30b-3p or NC was injected into TA muscle regeneration model every four days. Then, regenerating TA samples were harvested at days 3 and days 10, respectively. The staining on TA cross-sections revealed that overexpression of miR-30b-3p lead to less efficient regeneration characterized by the smaller myofiber cross-sectional area and more inflammatory cells, which is consistent with directly visible TA muscle injury status (Figure 5B-5D). The protein and mRNA of myogenic key factors were dramatically down-regulated in TA muscles administrated with agomiR-30b-3p at day 3 (Figure 5E-5N). These results indicate that superfluous miR-30b-3p is detrimental for muscle regeneration. In support of this notion, at 10 days after injection, TA muscle transfected by agomiR-30b-3p showed less efficient regeneration than NC according to the number and size of myofibers (Figure 6A-6M). Together, above evidence confirms that miR-30b-3p does regulate muscle development and regeneration of injury muscle.

**Figure 5.**
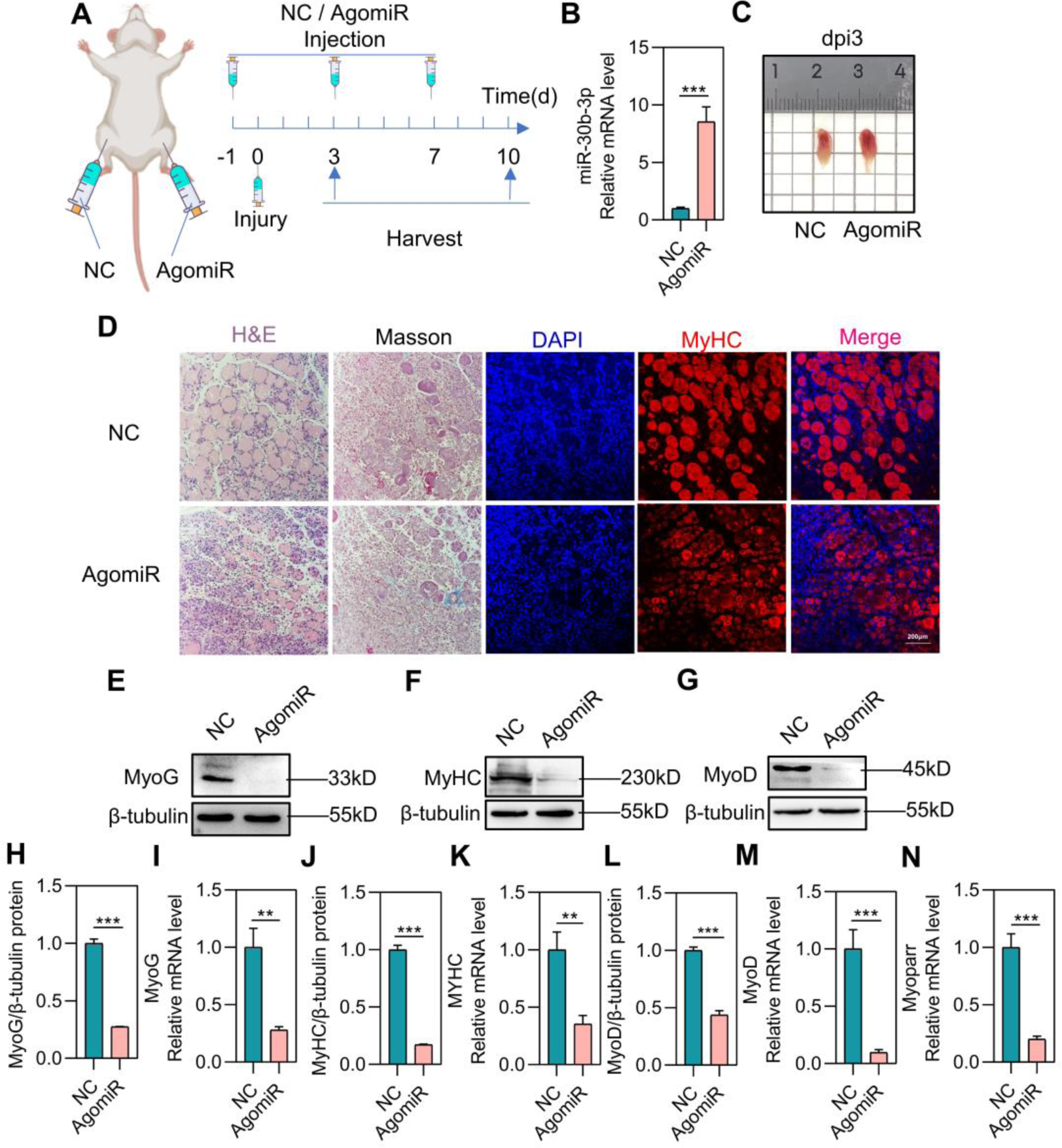
Elevated miR-30b-3p delays skeletal muscle regeneration (3dpi) (A) A schematic outlining the experimental protocol followed to analyze muscle regeneration. CTX was injected into the TA muscles and when the TA muscles were treated with CTX, it was defined as day 0. The left and right TA muscles were treated with NC and agomiR-30b-3p respectively, then harvested and analyzed at 3 days and 10 days post-injury. (B) Overexpression efficiency of agomiR-30b-3p targeting miR-30b-3p was detected from the extracted RNA of TA through qPCR after 3 days injury (n=3). (C) Representative images of regenerating TA muscles treated with NC and agomiR-30b-3p at day 3. (D) Immunofluorescence staining of MyHC, H&E staining and Masson trichrome staining were performed on cross sections of TA muscles after 3 days injury. (E-N) Western blot and qPCR detection of myogenic genes expression in the regenerating TA muscles at day 3 (n=3). Data are represented as mean ± SD. \**P* < 0.05; \*\**P* < 0.01; \*\*\**P* < 0.001 (Student’s *t* test).

**Figure 6.**
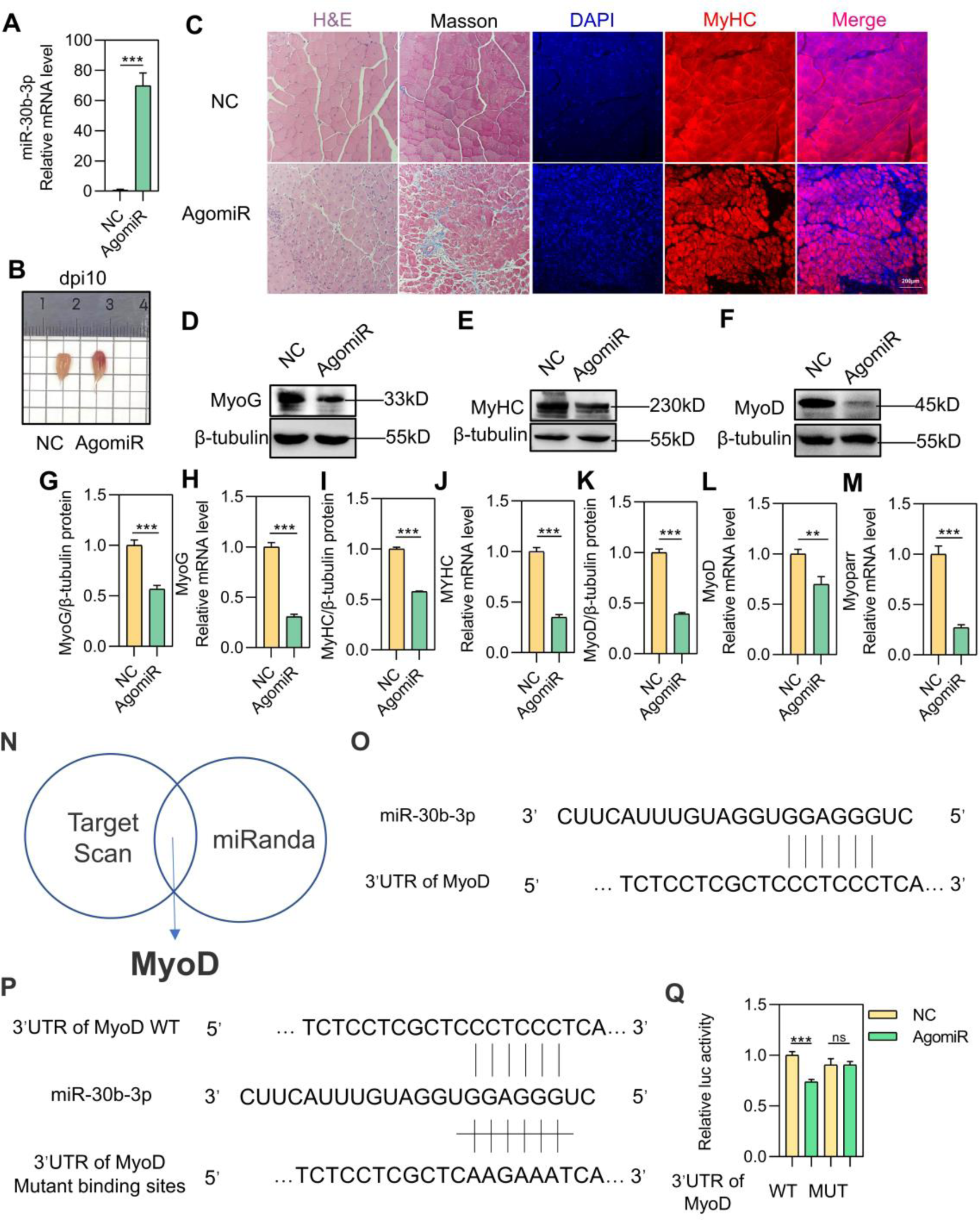
Elevated miR-30b-3p delays skeletal muscle regeneration (10dpi) and prediction and validation of MyoD 3’UTR as a binding target of miR-30b-3p. (A) Overexpression efficiency of agomiR-30b-3p targeting miR-30b-3p was detected from the extracted RNA of TA through qPCR after 10 days injury (n=3). (B) Representative images of regenerating TA muscles treated with NC and agomiR-30b-3p at day 10. (C) Immunofluorescence staining of MyHC, H&E staining and Masson trichrome staining were performed on cross sections of TA muscles after 10 days injury. (D-M) Western blot and qPCR detection of myogenic genes expression in the regenerating TA muscles at day 10 (n=3). (N) The binding target of miR-30b-3p predicted by TargetScan and miRanda was intersected. MyoD was selected for closely related to the myogenesis process. (O) Details of the alignment of miR-30b-3p to the binding site of MyoD 3’UTR. (P) Schematic of the double luciferase assay vector with upper wildtype (WT), and lower mutant (Mut) MyoD 3’UTR binding sites. (Q) AgomiR-30b-3p or NC were co-transfected with psiCHECK2-MyoD 3’UTR WT or psiCHECK2-MyoD 3’UTR Mut vectors into 293T cells. After 48 h, Dual-Luciferase reporter assay was quantified and normalized (n=4). Data are represented as mean ±SD. \**P* < 0.05; \*\**P* < 0.01; \*\*\**P* < 0.001 (Student’s *t* test).

### MiR-30b-3p targets the 3’UTR of MyoD

Given that miR-30b-3p makes a difference in myogenic differentiation, we further explored the molecular mechanisms of miR-30b-3p in myogenic regulation. In general, miRNAs are usually involved in the post-transcriptional level and bind to 3’ UTR of their target mRNA to suppress expression. Interestingly, MyoD was predicted to has a miR-30b-3p binding site in its 3’ UTR by both TargetScan and miRanda (Figure 6N-6O). Then the luciferase reporter vectors were constructed containing wild-type binding site of 3’ UTR (MyoD 3’ UTR-WT) or mutated binding site of 3’ UTR (MyoD 3’ UTR-MUT) (Figure 6P). As a result, only the luciferase activity of MyoD 3’ UTR-WT but not MyoD 3’ UTR-MUT could be reduced by agomiR-30b-3p (Figure 6Q).

### SNP rs3471653254 C>T regulates the transcription of Myogenin via affecting the binding of HOXA5

Above results have shown that SNPs in the upstream of MyoG affected myoblasts differentiation, but we wondered how the SNPs play the regulatory role. Therefore, based on the sequence of the promoter, we predicted SNP rs3471653254 C>T, which located at 299 sites before the TSS, is an important site for HOXA5 binding using the JASPAR database. We found HOXA5 keeps evolutionally conserved in multiple species and its binding site motif (Figure 7A-7B). To verify this speculation, the promoter segments harboring different SNPs were cloned into luciferase reporter vector (Figure 7C-7D). Reporter vectors were co-transfected into 293T cells with empty or plasmids expressing HOXA5. As illustrated in Figure 7E, HOXA5 obviously elevated the activity of promoters which contain the first SNP keeping T base. Moreover, the interaction between the promoter region and HOXA5 was confirmed by chromatin immunoprecipitation. Significantly higher HOXA5 enrichment levels were detected in promoter segment of DR than that of GS (Figure 7F). The mRNA and protein levels of HOXA5 were detected to authenticate the efficiency of overexpression. (Figure 7G-7I). Along with the up-regulation of HOXA5 on the promoter activity, MyoG and Myoparr expressing levels were significantly increased by 30%-70% in C2C12 cells (Figure 7J-7M). Overall, our findings elucidate that upstream binding site in which SNP rs3471653254 C>T exists is an enhancer of MyoG and Myoparr, and HOXA5 promotes their expression by being recruited.

**Figure 7.**
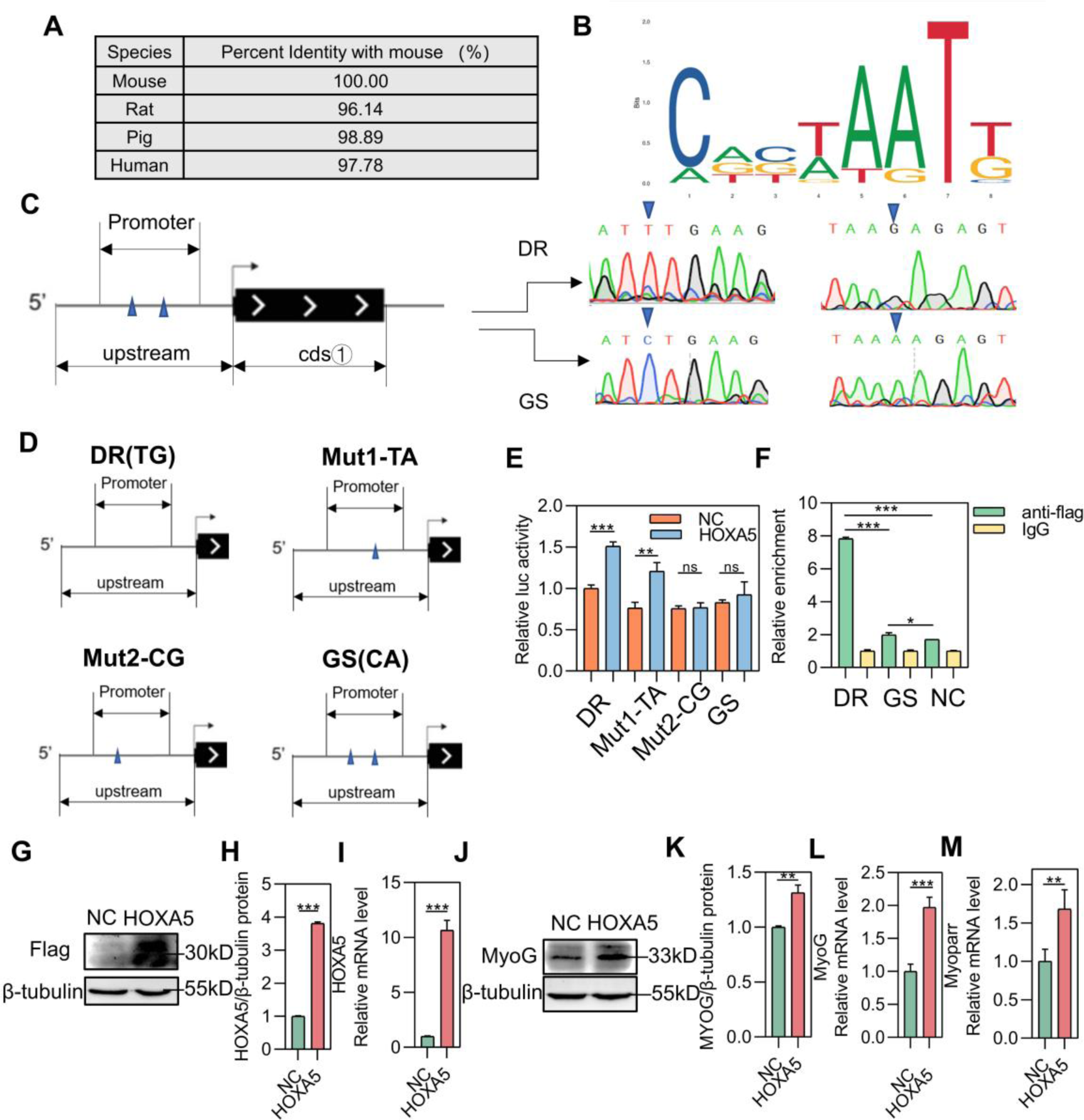
SNP rs3471653254 C>T regulates directly the interaction of HOXA5 and the promoter region. (A) The sequence similarity of HOXA5 was compared across four species. (B) Schematic of the possibility on HOXA5 combination motif. (C) The mutation loci information of the two pig breeds obtained by sequencing. (D) WT or two SNPs mutated in various combinations of promoter binding sites were inserted into reporter vectors. (E) HOXA5 overexpressed plasmid or NC were co-transfected with four above-mentioned reporter vectors from D into 293T cells. After 48 h, Dual-Luciferase reporter assay was quantified and normalized (n=4). (F) ChIP-qPCR analysis of the binding of HOXA5 in 293T cells transfected with two kinds of promoter segment vectors and NC. (G-I) The efficiency of HOXA5 overexpression compared with that of the NC detected by Western blot and qPCR. (J-M) Western blot and qPCR detection of MyoG and Myoparr expression in C2C12 cells transfected with HOXA5 overexpression vectors. Data are represented as mean ±SD. \**P* < 0.05; \*\**P* < 0.01; \*\*\**P* < 0.001 (Student’s *t* test).

## Discussion

The domestic pig is an essential protein source for humans. Genetic and phenotypic differentiation between pig breeds result from geographical divergence, local adaptation, and artificial selection(Ai et al. 2015; Li et al. 2017). Compared to Chinese indigenous pig breeds, western lean-type pig breeds have been subject to decades of artificial selection aimed at generating faster growth and higher muscle mass(Zhao et al. 2011; Li et al. 2017). These differences make them ideal research models to elucidate the potential mechanisms of phenotypic differentiation of muscle mass between pig breeds(Tang et al. 2007; Zhao et al. 2011; Zhao et al. 2015). Large-scale whole-genome resequencing revealed that SNP at MyoG region in pigs may contribute to the inter-breed differences(Wang 2021). Thereby, overexpression plasmids carrying different MyoG genotype were constructed respectively, and the MyoG plasmid of DR has a stronger promoting effect on myogenic differentiation. These findings are consistent with research showing the higher meat production and growth rate of western lean-type pig breeds, confirming the rationality and reliability of our results. As the MyoG genotype of DR is more conducive to the differentiation of myoblasts, we would like to explore which SNP play a major regulatory role. Through a series of experiments, we found that two SNPs in the upstream may have an important impact on myogenesis. Moreover, it also partially explains the differences in muscle mass between the two kinds of pig breeds.

Meanwhile, the promoter not only transcripts MyoG, but also transcripts lncRNA Myoparr, which is a recently discovered regulator of myogenesis in mouse and human transcribed from the upstream region of myogenin(Hitachi et al. 2019). Specifically high expression of Myoparr in skeletal muscles is consistent with its important roles in skeletal muscle. To unravel the underlying regulatory mechanism, we predicted the possible miRNAs that may be sponged by Myoparr and found miR-30b-3p, which has been previously proved to keep evolutionally conserved in multiple species (Figure S2B). In previous studies, miR-30b-3p functions in anticancer treatment by targeting multiple genes(Gao et al. 2019; Li et al. 2020; Chen et al. 2022). Here, when miR-30b-3p was overexpressed, the expression of Myoparr WT was obviously inhibited, but fluorescence activity of Myoparr MUT was not affected. In accordance with our expectation, miR-30b-3p holds an opposite expressing tendency to Myoparr no matter during the differentiation stage of C2C12 or during embryonic muscle development. To explore the precise effect of miR-30b-3p on skeletal muscle, we explore its function by employing overexpression and knockdown strategy. It was found that miR-30b-3p not only prevents myoblast differentiation, but also delays the muscle regeneration. From the above, the conclusion can be reached that Myoparr forms a ceRNA regulatory network with miR-30b-3p, and regulates the myoblast differentiation together. In addition, the function and mechanism of Myoparr-miR-30b-3p ceRNA in myogenesis are conserved among pigs and mice, which suggests that Myoparr-miR-30b-3p ceRNA have more influence on muscle development.

Increasing evidence has demonstrated that miRNAs were involved in the muscle development via interacting with 3’UTR of target mRNAs leading to mRNA degradation or translational repression(Yang et al. 2021). MyoD, the key myogenic differentiation factor, has been well known as a master TF in myogenic cell-lineage specification during development and trans-differentiation(Buckingham and Tajbakhsh 1999). What’s more, MyoD is a 3D genome structure organizer for muscle cell identity(Lassar et al. 1986; Davis et al. 1987; Tapscott et al. 1988; Weintraub et al. 1989; Rudnicki et al. 1993). In this study, it was confirmed as the target of miR-30b-3p which is a novel myogenic inhibitor.

Promoters are specific DNA sequences that RNA polymerase II binds to and initiate transcription. RNA polymerase II requires transcription factors to be recruited to the transcription start site as part of a large transcription pre-initiation complex(Harris and Marles-Wright 2019). The high conservatism of HOXA5 suggests that it serves a vital function. More importantly, HOXA5 is a crucial transcriptional factor in both tumor suppression and oncogenesis via the interaction with various target genes and signaling pathways(Liao et al. 2020; Fan et al. 2022; He et al. 2022; Jin et al. 2023). Once SNP rs3471653254 is changed from T to C, the enrichment level of HOXA5 to the promoter is down-regulated, which subsequently inhibits the promoter activity of MyoG and Myoparr. As described, HOXA5 as a transcription factor activates the transcription of MyoG and Myoparr.

Taken together, our research elucidates the regulatory network of SNP rs3471653254 in two kinds of pig breeds and a new myogenic inhibitor miR-30b-3p as detailed in Figure 8. Activation of RNA polymerase II transcription in the promoter region shared by MyoG and Myoparr requires the recruitment of HOXA5. In western lean-type pig breeds, SNP rs3471653254 base presented as T, in which a large amount of HOXA5 could be enriched, thus promoting the transcription activity of Myoparr and MyoG. However, in Chinese indigenous pig breeds, the enrichment level of HOXA5 was much lower for the base C. Subsequently, Myoparr acts as a sponge of miR-30b-3p, so the expression of Myoparr will also influence the ability to sponge miR-30b-3p. Meanwhile miR-30b-3p inhibits the expression of MyoD through interacting with 3’UTR of MyoD, then down-regulating the progress of myoblast differentiation. Certainly, there is no denying that SNP rs3471653254 C>T could affect myoblast differentiation through this pathway that regulating MyoG transcription. More importantly, we revealed a crucial Myoparr/mir-30b-3p/MyoD axis for myogenesis. Future studies using comprehensive analysis of other interacting factors will further define the roles of Myoparr/mir-30b-3p/MyoD axis in skeletal muscle formation and disorders affecting muscles. Thus, our findings suggested that the regulation of Myoparr/mir-30b-3p/MyoD axis may be a useful therapeutic strategy for myopathy. Our study provides a theoretical basis for explaining the differences in meat production between western lean-type pig breeds and Chinese indigenous pig breeds. Moreover, this SNP could be used for improving meat production in swine industry in the future. These mechanisms can now be interpreted in molecular detail and provide an exciting theoretical framework for past and future functional studies of SNP, giving us unique insight into the complex process of species evolution.

**Figure 8.**
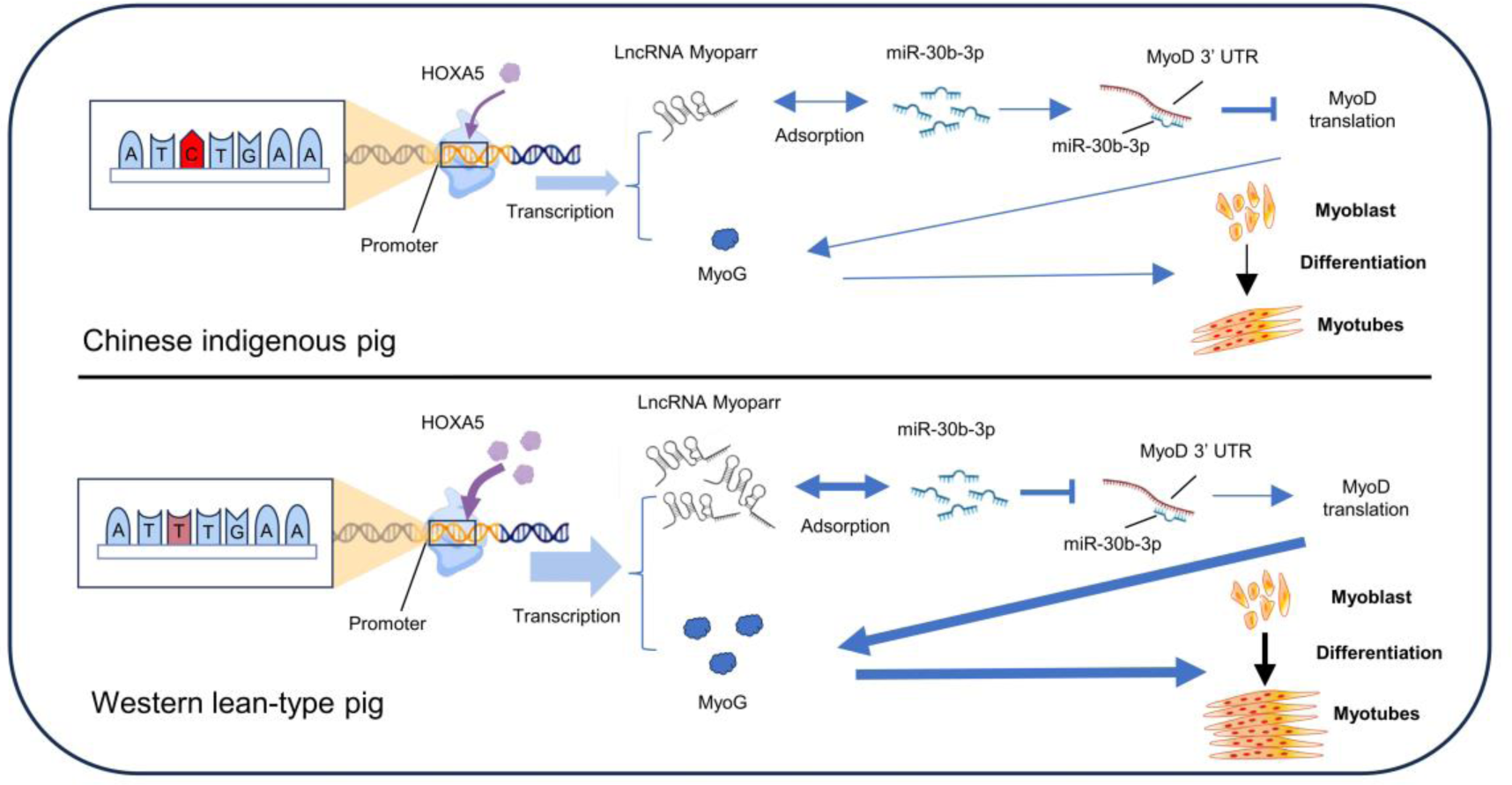
SNP rs3471653254 C>T associated with lncRNA Myoparr sponging miR-30b-3p regulates myogenic differentiation via affecting the binding transcription factor HOXA5. Schematic diagram of the mechanism by which SNP rs3471653254 C>T modulates myogenesis. SNP rs3471653254 C>T influences the enrichment of HOXA5 to regulate the transcriptional expression level of MyoG and Myoparr. Myoparr can be used as the sponge of mir-30b-3p for adsorption. Afterwards miR-30b-3p inhibits the expression of MyoD through interaction 3’UTR of MyoD, then down-regulating the process of myoblast differentiation.

## Materials & methods

### Animal

Wild-type mice (C57BL/6) were purchased from Cyagen Biosciences Co., Ltd. (Suzhou) and were housed under specific-pathogen-free conditions. All mice used in this study had a C57BL/6J genetic background and were housed in SPF conditions during the experiment. Guangdong small-ear spotted and Duroc pigs were acquired from the Guangdong YIHAO Food Co. Ltd (Guangzhou, Guangdong Province). All animals were kept in an environment in which they had sufficient food, were allowed to move freely, and had suitable lighting. Housing, husbandry and all experimental protocols for mice and pigs were approved and performed in accordance with the guidelines established by the Animal Care and Use Committee of Guangdong Province and conducted according to ethical standards.

### Cardiotoxin injury

Cardiotoxin (CTX, Sigma) induced muscle injury/regeneration model in adult mouse is an established, reliable model to study muscle regeneration(Goetsch et al. 2003). CTX was dissolved in sterile saline to a final concentration of 10 mmol/L. The mice’s hindlimbs were cleaned with 75% alcohol. 50 µL of 10 mmol/L CTX was injected into the left and right TA muscles which have been transfected with miR-30b-3p agomir and NC one day before, respectively. Then regenerating TA muscles were harvested 3 and 10 days after CTX injection. These tissues were cleaned in phosphate buffer saline (PBS) and were quickly frozen in liquid nitrogen for future experiments.

### Primary myoblasts isolation and cell culture

Dorsal muscle of porcine E35 embryo was isolated and separated by 1 mg/ml type I collagenase (Sigma) in Dulbecco’s modified Eagle’s Medium (DMEM, Gibco) at 37℃ for 1.5-2 h. The suspension was fermented with ten milliliters of culture media (20%

FBS/DMEM) and centrifuged at 1600 × g for 10 min. The pellet was resuspended in 10 ml of growth media (20% FBS/DMEM + 2.5ng/ml bFGF), filtered through a 100 µm cell strainer, and plated on a 10 cm Matrigel coated culture dish. The culture is incubated in a small volume of PBS, gently tapping the plate to displace the myoblasts to enrich the myoblasts. When cells growing into confluence, the culture medium was replaced to DMEM with 2% horse serum to induce differentiation.

The C2C12 mouse myoblast cell line was supplied by American Type Culture Collection (ATCC). C2C12 myoblasts were cultured in DMEM containing 10% foetal bovine serum and 1% penicillin/streptomycin in a humidified atmosphere with 5% CO2 at 37 °C. When cells reached 90% confluence, the medium was replaced into DMEM with 2% horse serum and 1% penicillin/streptomycin to induce differentiation.

### Overexpression

The sequences of MyoG and HOXA5 were cloned into pcDNA3.1 vector. C2C12 cells were seeded into 6-or12-well plates at 12 hours before treatment and then transfected with expression plasmids using MK40 (MIKX) according to the manufacture’s instruction. Transfections were performed at least in triplicate for each experiment.

### MiR-30b-3p agomir and antagomir treatment

MiR-30b-3p agomir and antagomir (RiboBio), were used to specifically up-regulate or down-regulate miR-30b-3p activity. Following the instructions, agomir and antagomir dissolved to a concentration of 20µM were added directly into the culture medium of cells. After transfection for 12 h, the culture medium was substituted by fresh DMEM containing 10% FBS. After 2 days of treatment, cells were harvest to detect the efficiency of activation or suppression.

### RNA extraction and real-time quantitative PCR (qPCR)

The cells and regenerating tibialis anterior (TA) muscles were washed with PBS and harvested for total RNA extraction using TRIzol reagent (vazyme). 1 μg of the total RNA was processed into single-stranded complementary DNA (cDNA) using a StarScript II First-strand cDNA Synthesis Kit (GenStar) following manufacturer’s instructions. Quantitation of the mRNA levels by qPCR was performed on a real-time PCR system using SYBR Green Master Mix (GenStar). The relative mRNA expression level was normalized to that of GAPDH. Gene expression was quantified by comparative CT method. The primers used for qPCR are listed in Table 1.

### Western blot

Protein extracts of cultured C2C12 cells or TA muscles were obtained using protein extraction buffer (FDbio) within 1% protease inhibitor cocktail. The lysed samples were centrifuged at 14, 000g and 4°C for 10 min and supernatants containing protein were collected. Then 5X protein loading buffer was added to the lysates prior to their full denaturation in boiling water for 10 min. Equivalent amount of protein samples were separated in 8% or 10% SDS-PAGE and transferred to polyvinylidene difluoride membrane (Millipore). After blocked with 4% bovine serum albumin (BSA) for 1 hour, membranes were incubated overnight at 4°C with the specific primary antibodies shown in Table 2. The membranes were washed with Tris-buffered saline with Tween 20 (TBST) and then incubated with corresponding secondary antibodies for 1 h at room temperature. Blots were visualized by the ECL chemiluminescence system.

### Immunofluorescence

The cells cultured in 12-well plates or 24-well plates were fixed in 4% paraformaldehyde for 10 minutes, followed by permeabilization in 0.5% Triton X-100 in PBS for 15 minutes. Samples were then blocked in 4% bovine serum albumin in PBS for 1 h at RT. Then, the cells were incubated with primary antibodies overnight at 4°C. Washed three times with PBS, the cells were incubated with secondary antibodies for 1 hour at room temperature. Finally, the cells were washed thrice in PBS, and nuclei were stained by DAPI. Antibodies are listed in Table 2. Immunostaining images were obtained via fluorescent reverse microscopy (Nikon).

### Histology

Freshly isolated adult TA muscles were immediately fixed in 4% paraformaldehyde at 4 °C for 18 h and subsequently embedded in paraffin. 4 μm thick cross-sections of TA muscles were subjected to H&E staining and masson’s trichrome staining which was performed according to procedures provided by the Hematoxylin-eosin (H&E) staining kit and masson’s Trichrome Stain Kit.

### Luciferase reporter assay

Luciferase reporter plasmids (psiCHECK2, sangon) were transfected into 293T cells seeded in a 24-well plate. After 36 h transfection, cells were lysed and enzymic reactions were assayed by using the Dual Luciferase reporter assay system (Promega). The firefly luciferase activity was normalized to renilla luciferase internal control to exclude the differences of transfection efficiency.

### Chromatin immunoprecipitation

293T cells transfected were cross-linked with 1% formaldehyde for 10 minutes at room temperature. The fixing solution was quenched for 5 min by adding glycine. Cells were lysed in lysis buffer (50 mM Tris-HCl pH 8.0, 10 mM EDTA, 0.5% SDS, 20 µg/ml proteinase K) on ice and chromatin was sonicated using a Covaris Sonicator for 8 min to generate chromatin fragments of 200-300 bp DNA. The clarified nuclear extracts were incubated with IgG antibody applied as negative control or target antibody shown in Table 2 with rotation overnight at 4°C and conjugated with Chromatin immunoprecipitation-grade protein G magnetic beads (Cell Signaling Technology). After extensive washing, bound DNA fragments were purified and eluted by elution buffer. The enrichment of DNA sequences was analyzed via qPCR using specific primers. Data were normalized to the respective control IgG values and assessed relative to the input DNA.

### Statistical analysis

All dot plots and graphs were generated using the Prism 9 software (GraphPad Software). Experiments were done with a minimum of three biological replicates. Data are presented as mean ± SEM, and the statistical significance analysis was performed using an unpaired two-tailed Student’s t test to test differences between groups. The level of significance is indicated as follows: * p < 0.05; ** p < 0.01; *** p < 0.001.

## Acknowledgments

This work was supported by the National key research and development program (2023YFD1300201), National Natural Science Foundation of China (32072697), Selection and Breeding of New Local Pig Breeds and Promotion of Industrialization (2022-440000-43010101-9501), Selection and Breeding of Guangdong small-ear spotted pig (2022-440000-4301030202-9510). To acknowledge all of the people who have contributed to this paper.

## Author Contribution

Delin Mo: Conceptualization, Resources, Supervision, Methodology, Funding acquisition, Writing – review & editing. Zhuhu Lin: Methodology, Formal analysis, Software, Validation, Investigation, Visualization, Writing – original draft. Xiaoyu Wang: Methodology, Formal analysis. Ziyun Liang: Project administration. Rong Xu: Data curation. Meilin Chen: Data curation. Xian Tong: Data curation. Chenggan Li: Data curation. Renqiang Yuan: Project administration. Yaosheng Chen: Resources. Xiaohong Liu and Yunxiang Zhao: Funding acquisition.

## Conflict of Interest Statement

The authors declare no competing interests.

## Ethics Statement

The animal experimental procedures used in this experiment were approved by the Animal Care and Use Committee of Guangdong Province, China. Approval ID or permit numbers are SCXK (Guangdong) 2011-0029 and SYXK (Guangdong) 2011-0112.

## Data Availability Statement

All data that support the findings of this study are available from the corresponding author upon reasonable request.

## Declaration of interest statement

The authors declare no competing interests.

